# Bulk and spatially resolved extracellular metabolomics of free-living nitrogen fixation

**DOI:** 10.1101/2022.01.13.476280

**Authors:** Darian N Smercina, Young-Mo Kim, Mary S Lipton, Dusan Velickovic, Kirsten S Hofmockel

## Abstract

Soil microorganisms drive ecosystem function, but challenges of scale between microbe and ecosystem hinder our ability to accurately quantify and predictively model the soil microbe-ecosystem function relationship. Quantifying this relationship necessitates studies that systematically characterize multi-omics of soil microorganisms and their activity across sampling scales from spatially resolved to bulk measures, and structural complexity, from liquid pure culture to *in situ*. To address this need, we cultured two diazotrophic bacteria in liquid and solid media, with and without nitrogen (N) to quantify differences in extracellular metabolites associated with nitrogen fixation under increasing environmental structural complexity. We also quantified extracellular metabolites across sampling scales including bulk sampling via GC-MS analysis and spatially resolved analysis via MALDI mass spectrometry imaging. We found extracellular production of inorganic and organic N during free-living nitrogen fixation activity, highlighting a key mechanism of terrestrial N contributions from this process. Additionally, our results emphasize the need to consider the structural complexity of the environment and spatial scale when quantifying microbial activity. We found differences in metabolite profiles between culture conditions, supporting previous work indicating environmental structure influences microbial function, and across scales, underscoring the need to quantify microbial scale conditions to accurately interpret microbial function.

**Importance:** Studying soil microorganisms, both who is present and what they are doing, is a challenge because of vast differences in scale between microorganism and ecosystem and because of inherent complexities of the soil system (e.g., opacity, chemical complexity). This makes measuring and predicting important ecosystem processes driven by soil microorganisms, like free-living nitrogen fixation, difficult. Free-living nitrogen fixing bacteria play a key role in terrestrial nitrogen contributions and may represent a significant, yet overlooked, nitrogen source in agricultural systems like bioenergy crops. However, we still know very little about how free-living nitrogen fixation contributes nitrogen to terrestrial systems. Our work provides key insight by hierarchically increasing structural complexity (liquid vs. solid culture) and scale (spatially resolved vs. bulk) to address the impact of environmental structure and sampling scale on detection of free-living nitrogen fixation and to identify the forms of nitrogen contributed to terrestrial systems by free-living nitrogen bacteria.

## Introduction

Soil microorganisms are a key link between above and belowground ecosystem function, driving energy and nutrient transfer between the atmosphere, biosphere, and pedosphere (1–3). Multi-omic analysis aimed at understanding the structure and function of these microorganisms has become a routine tool for environmental samples. Despite generating large amounts of data, even quantitatively linking ‘omics of a specific function (e.g. functional genes and proteins) to measures of that function is often unsuccessful (2–6) because of inherent challenges of studying soils (5–7) and soil microorganisms. Given these challenges, the importance of linking microbial community structure to function has been questioned (8, 9), however without clearer understanding of how microorganisms relate to observed functions we cannot yet rule out the importance of individual microbial community members. Our limited ability to quantitively link soil microbial communities to ecosystem function hinders our understanding of ecosystem processes, leaving us vulnerable to losing vital ecosystem services provided by our soils, particularly in the face of climate change (10).

Quantifying the link between soil microorganisms and ecosystem function requires systematic studies that characterize multi-omics of soil microorganisms and their functions hierarchically across scales of space and complexity (Fig. 1; (4, 11). *In vitro* studies using pure cultures or limited species are an appealing option and have the potential to provide fundamental microbial and ecological knowledge (10, 12, 13). However, culturing conditions are often quite different from those experienced by microorganisms in soil and attachment to surfaces has been shown to impact microbial growth and function (14, 15). Thus, physical structure influences microbial function, and it is therefore essential for studies to systematically characterize multiomics and function *in vitro* under growth conditions of increasing complexity (e.g. environmental structure) in order to determine how culture work may better inform *in situ* processes (Fig. 1).

**Fig. 1:**
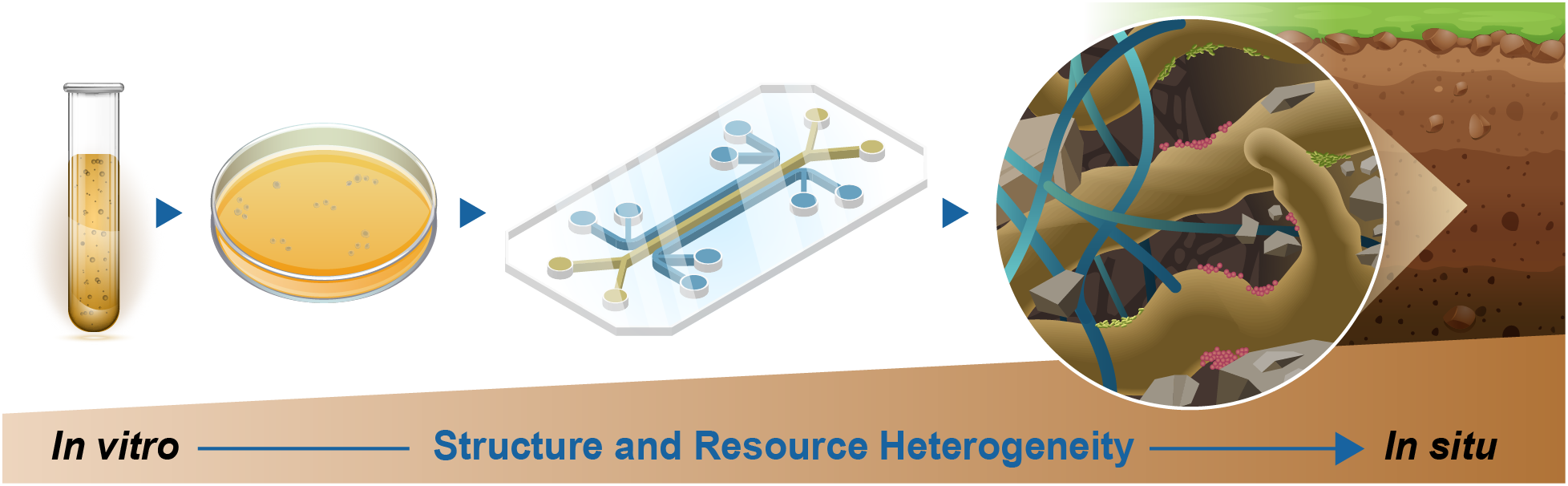
Depiction of the systematic scaling of system complexity from liquid and solid culture to synthetic soils (represented here as a microfluidic chip) to *in situ* soil conditions. In our study we focus on the first two steps, relating metabolomics in liquid and solid media culture.

In this study, we explored two questions surrounding the microbe-ecosystem function relationship: (1) Are there microbial metabolomic signatures associated with target ecosystem processes? (2) How do growth conditions and sampling scale influence metabolomic signatures? We used free-living nitrogen fixation (FLNF), biological nitrogen fixation (BNF) carried out by heterotrophic bacteria (diazotrophs), as a model microbial process and ecosystem function to address these questions. FLNF is carried out by a wide diversity of soil bacteria and occurs in all terrestrial biomes, contributing significantly to terrestrial N (16, 17). These contributions are thought to occur predominately through release of ammonium and organic N sources, like amino acids, as occurs during symbiotic BNF (18–21). Thus, extracellular metabolites may be valuable to understanding FLNF and identifying biosignatures.

We examined extracellular metabolites from two diazotrophic bacteria cultured under conditions that promote (N-free) or inhibit (N-rich) FLNF. Additionally, cultures were grown in liquid or on solid media to examine the impact of physical structure on detected metabolites. Lastly, we examined extracellular metabolites across sampling scale from spatially resolved measures using matrix-assisted laser desorption/ionization mass spectrometry imaging (MALDI– MSI) to bulk sampling via gas chromatography–mass spectrometry (GC-MS) analysis. We hypothesized: (1) bulk metabolite profiles differ between N-rich and N-free conditions and also between liquid and solid cultures, (2) there are extracellular metabolites associated with FLNF activity and only observed under N-free conditions, (3) these extracellular metabolites are detectable with bulk and spatially resolved sampling, and (4) N-containing compounds are produced during FLNF and are readily detectable in bulk and spatially resolved extracellular metabolites under N-free conditions.

## Results

### Microbial biomass – total biomass, biomass C, and biomass N

Total microbial biomass, including cells and associated debris such as EPS, was collected from all treatments except for the AV N-rich solid treatment. In this case, microbial colonies had grown into and below the agar surface and it was not possible to collect biomass. Total biomass was highly variable across all treatments (CV across treatments ranged from 4.6% to 100.4%) and there were no significant differences observed with N treatment, culture type, or organism (Fig. 2D). Both biomass C and N differed significantly across treatments (Fig. 2A and 2B, respectively). In general, C and N content were greater in N-rich treatments where N was readily available compared to N-free treatments. There was also a trend towards greater biomass C and N from solid media, but this was mostly observed in the N-rich treatments. These C and N values translated to C:N ratios that predominately differed only between N treatments with the N-free treatment resulting in greater biomass C:N ratios than N-rich conditions, regardless of culture type or organism (Fig. 2C).

**Fig. 2:**
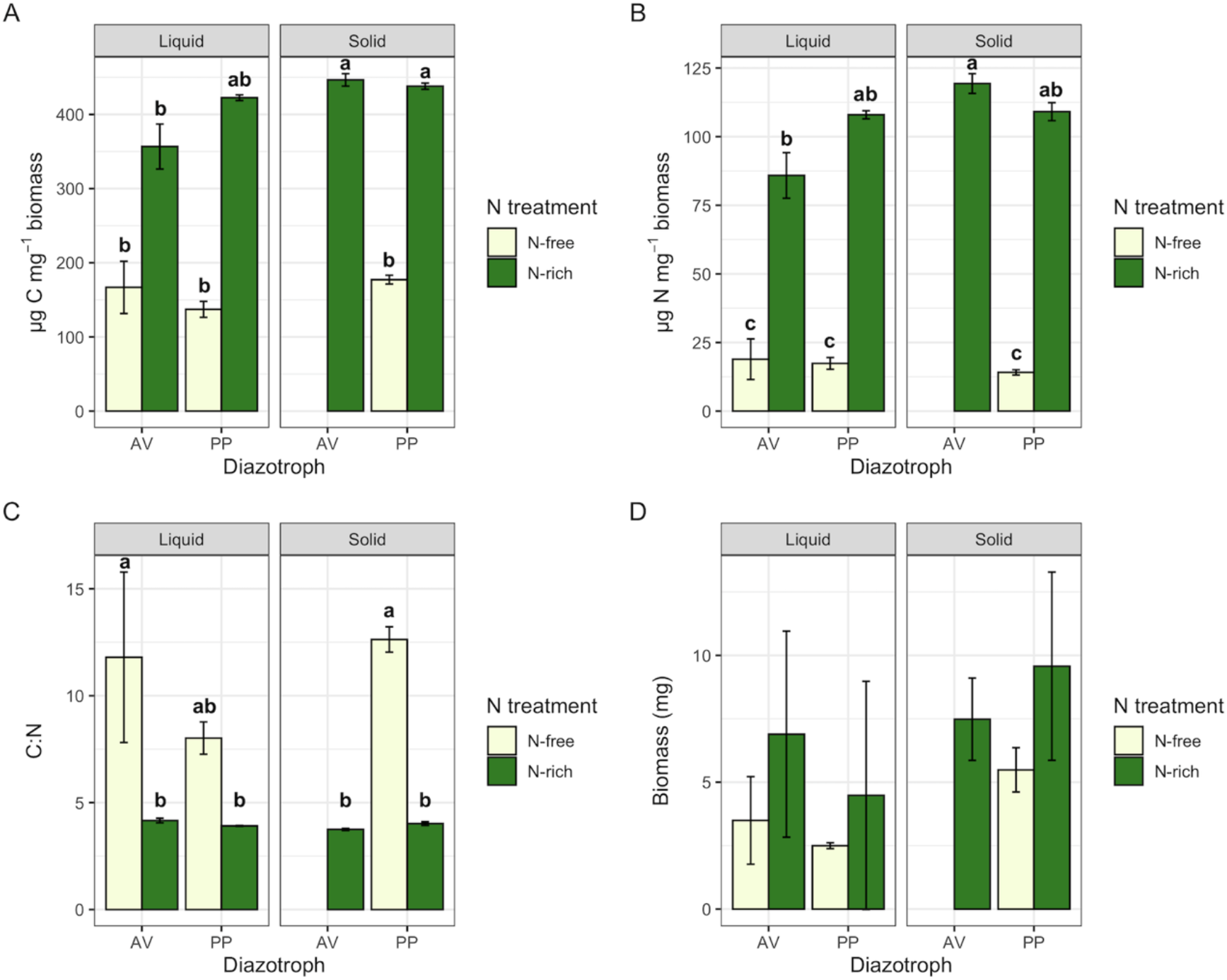
Microbial (A) C content, (B) N content, (C) C:N ratio, and (D) total biomass. Bars represent average values ± standard error and are colored by nitrogen treatment. Figures are faceted by culture type. Lowercase letters represent significant difference at p < 0.05.

### Extracellular ammonium availability

Extracellular ammonium availability was measured supernatant and rinsate samples and was detected in all treatments regardless of culture type, N treatment, or organism. On a per unit biomass basis, ammonium concentrations differed significantly by organism (F = 16.390, p = 0.0012) and by the interaction between culture type and N treatment (F = 35.411, p < 0.0001). Extracellular ammonium availability per unit biomass was over 8x greater in PP than AV cultures (Fig. 3A). Under N-free conditions, ammonium availability was greater in liquid than in solid culture while in N-rich conditions the opposite was observed (Fig. 3B). Extracellular ammonium availability is of particular interest in N-free treatments as ammonium is hypothesized to be released from cells actively fixing N and thus represents a major form of N contributed by FLNF to terrestrial systems (18, 33). Therefore, we also calculated the percent of fixed N available as extracellular ammonium (Fig. 4). Total fixed N was estimated as total biomass N measured in N-free treatment samples. We find only upwards of 7.5% (± 2.3) of fixed N is readily available as extracellular ammonium. This did not differ significantly between culture types (F = 2.171, p = 0.184), though ammonium concentrations in solid culture tended to be lower than those in liquid culture.

**Fig. 3:**
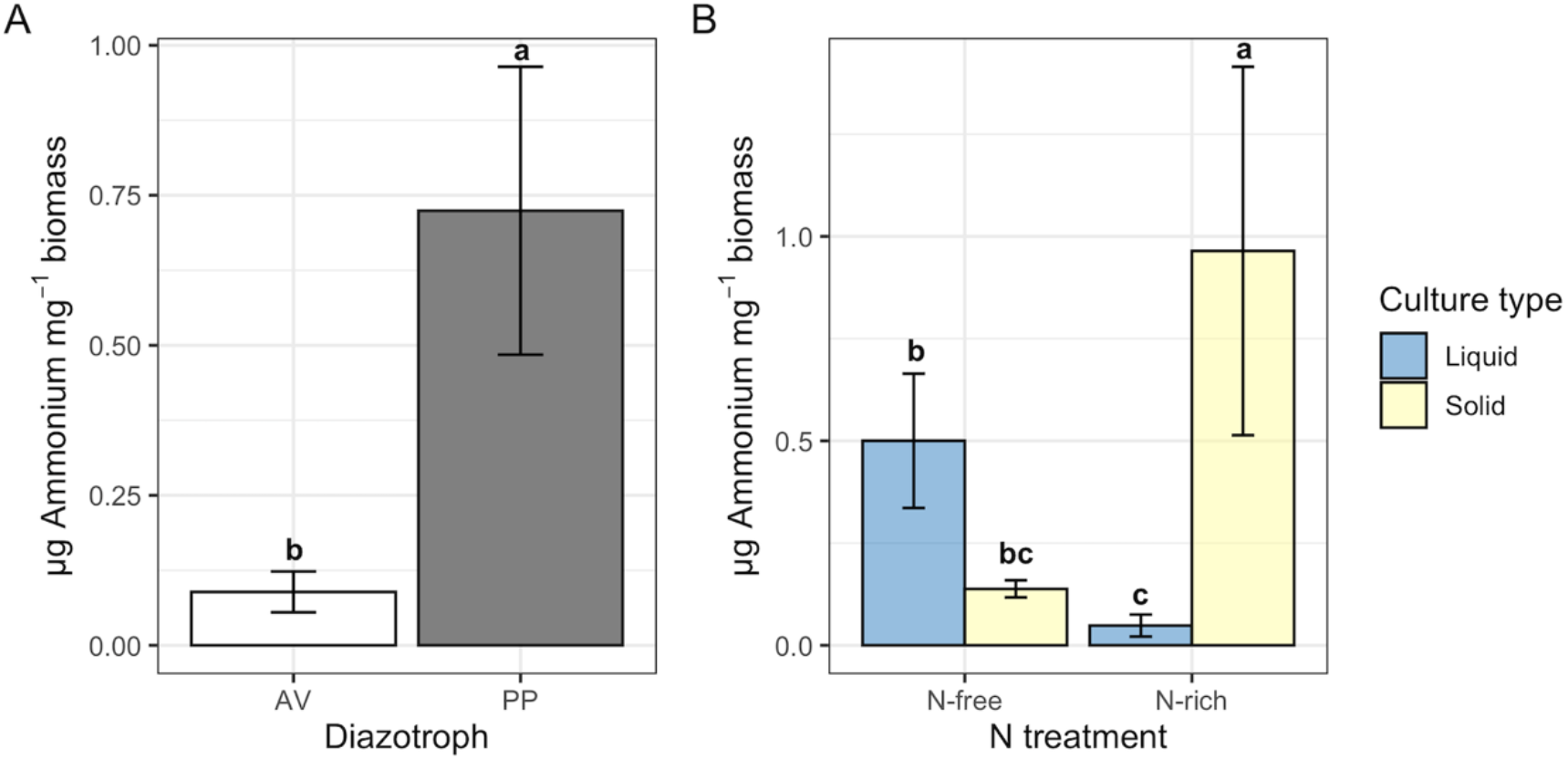
Extracellular ammonium availability per unit biomass shown by (A) diazotrophic organism and by (B) N treatment. Bars represent average values ± standard error. Bars in (B) are colored by culture type. Lowercase letters indicate significant difference at p < 0.05.

**Fig. 4:**
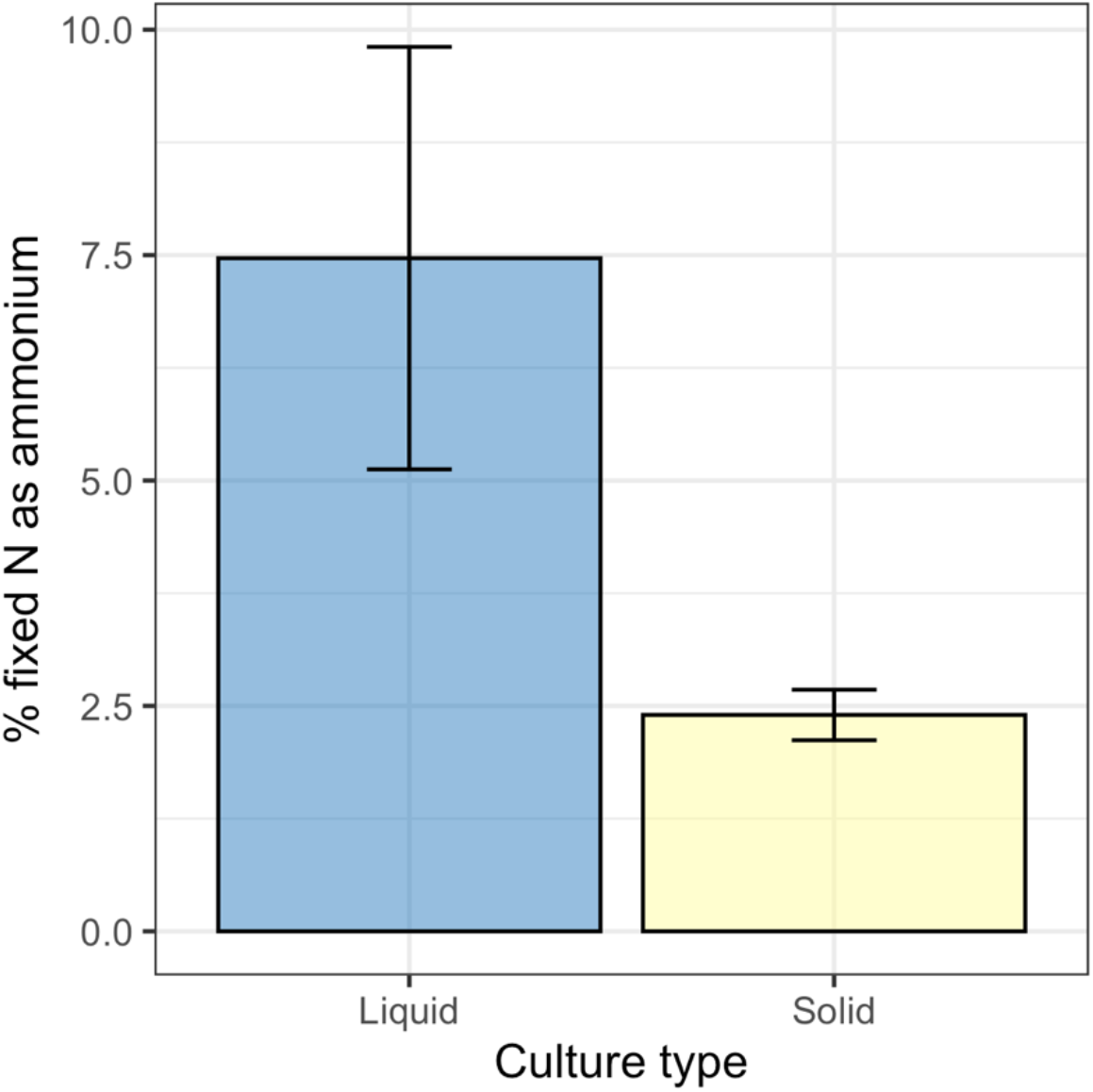
Percent of fixed nitrogen available as extracellular ammonium in N-free treatments. Bars represent average ± standard error. No significant difference was observed between culture types.

### Bulk extracellular metabolites

Across all treatments, 307 metabolites were detected with bulk sampling and of these 93 were successfully annotated (>80% confidence). The total number of detected metabolites differed between treatment groups (Supp. Fig. 2) with generally more metabolites detected in N-free treatments, the majority of which were within the unannotated portion of detected metabolites. Distinct metabolite profiles, represented by Bray-Curtis and Jaccard distance based on all detected metabolites, were observed between N treatment, culture type, and their interaction (Fig. 5; Table 1). N treatment and culture type, together with their interaction, explain over 50% of the variance in metabolite profiles based on peak intensity (Fig. 5A) and based on presence-absence (Fig. 5B). Metabolite profiles based on abundance separated predominantly by culture type and then by N treatment (Fig. 5A). Metabolite profiles based on presence-absence show clear separation between culture types for N-rich treatments but have little separation under N-free conditions (Fig. 5B).

**Table 1:**
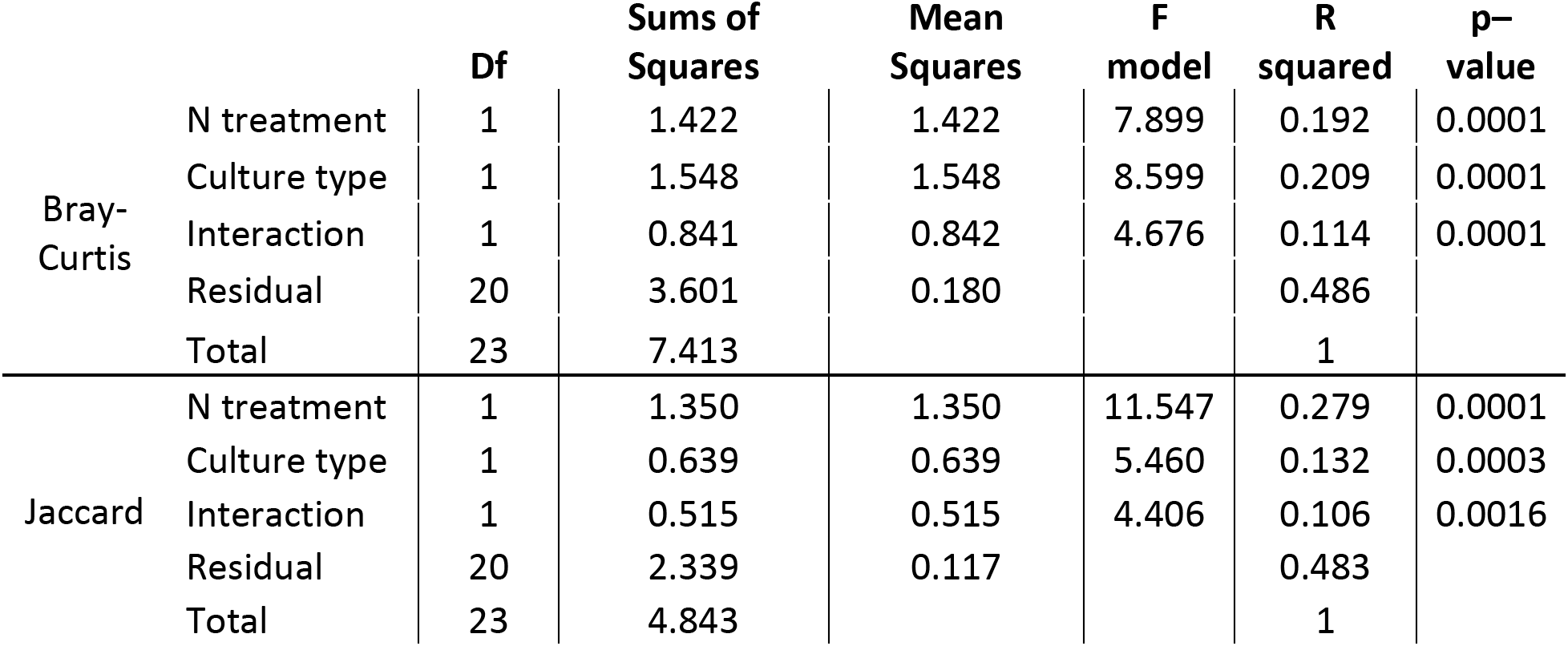
PERMANOVA results for Bray-Curtis and Jaccard distance of macroscale metabolite abundance.

|  |  | Df | Sums of Squares | Mean Squares | F model | R squared | p-value |
| --- | --- | --- | --- | --- | --- | --- | --- |
| Bray-Curtis | N treatment | 1 | 1.422 | 1.422 | 7.899 | 0.192 | 0.0001 |
|  | Culture type | 1 | 1.548 | 1.548 | 8.599 | 0.209 | 0.0001 |
|  | Interaction | 1 | 0.841 | 0.842 | 4.676 | 0.114 | 0.0001 |
|  | Residual | 20 | 3.601 | 0.180 |  | 0.486 |  |
|  | Total | 23 | 7.413 |  |  | 1 |  |
| Jaccard | N treatment | 1 | 1.350 | 1.350 | 11.547 | 0.279 | 0.0001 |
|  | Culture type | 1 | 0.639 | 0.639 | 5.460 | 0.132 | 0.0003 |
|  | Interaction | 1 | 0.515 | 0.515 | 4.406 | 0.106 | 0.0016 |
|  | Residual | 20 | 2.339 | 0.117 |  | 0.483 |  |
|  | Total | 23 | 4.843 |  |  | 1 |  |

**Fig. 5:**
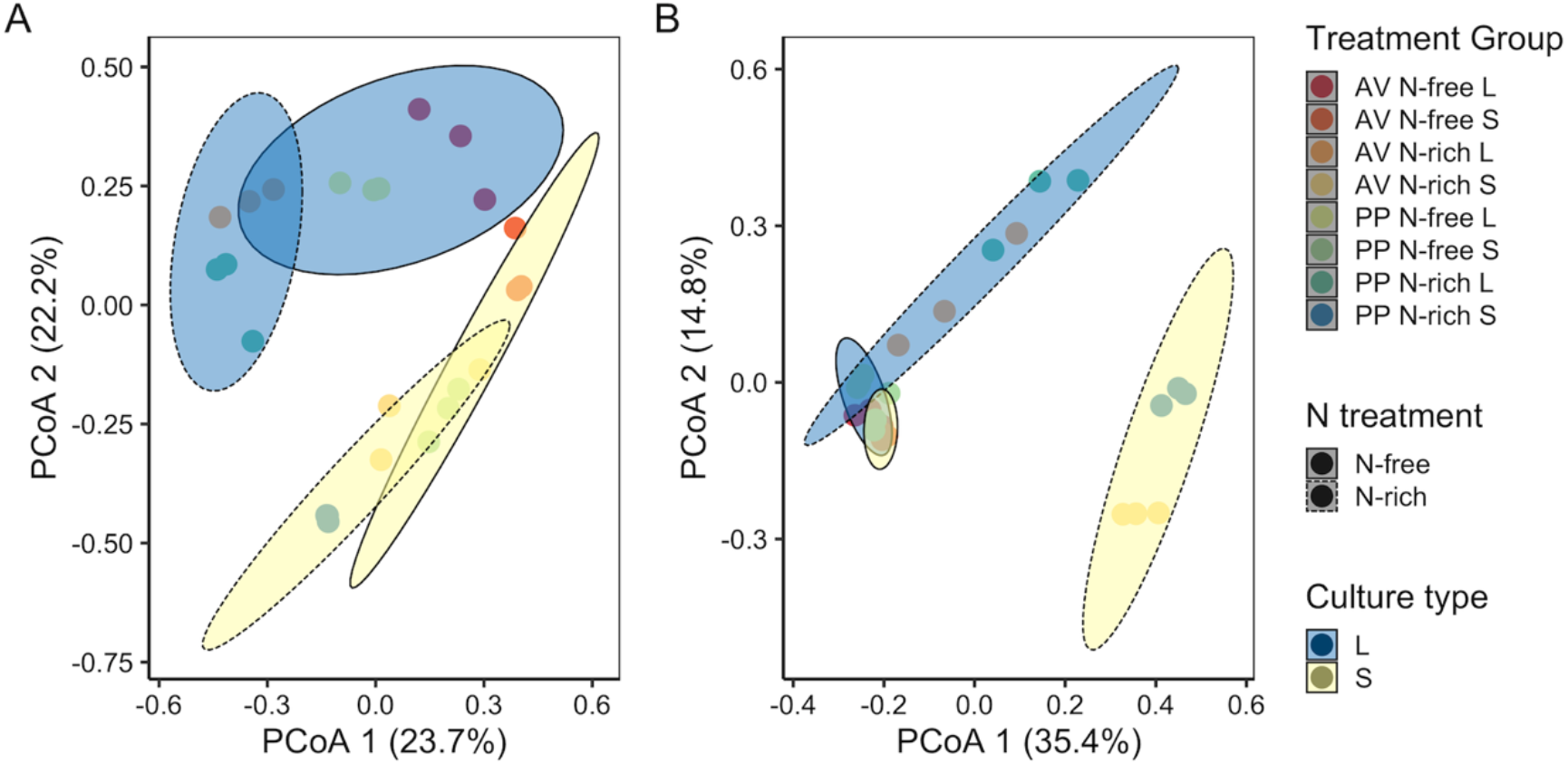
Principal coordinates analysis (PCoA) of metabolite chemistry based on (A) Bray-Curtis of peak intensity and (B) Jaccard of presence-absence including all detected metabolites. Each point represents a single sample and are colored by treatment group (organism, N treatment, culture type). 95% confidence ellipses are shown for culture type, represented by color, and N treatment, represented by line type.

Because FLNF activity is hypothesized to result in the release of N-containing metabolites (33), we focused on N-containing extracellular metabolites. Of the 93 annotated metabolites detected through bulk sampling, 35 were N-containing. We found significant differences in N-containing compounds across treatments, after correcting for background metabolites, with significant interactions between N treatment, culture type, and organism (F = 10.695, p = 0.0048). N-free treatments were richer in N-containing metabolites (Supp. Fig. 3) but had similar or significantly lower total abundances of N-containing metabolites compared to N-rich treatments (Fig. 6). Examining the specific composition of these N-containing compounds, we found a variety of amino acids in N-free samples not well represented in N-rich samples (Supp. Fig. 3), but only a few N-containing metabolites were unique to N-free conditions including pantothenic acid, L-pyroglutamic acid, L-glutamic acid, and 4-pyridoxic acid.

**Fig. 6:**
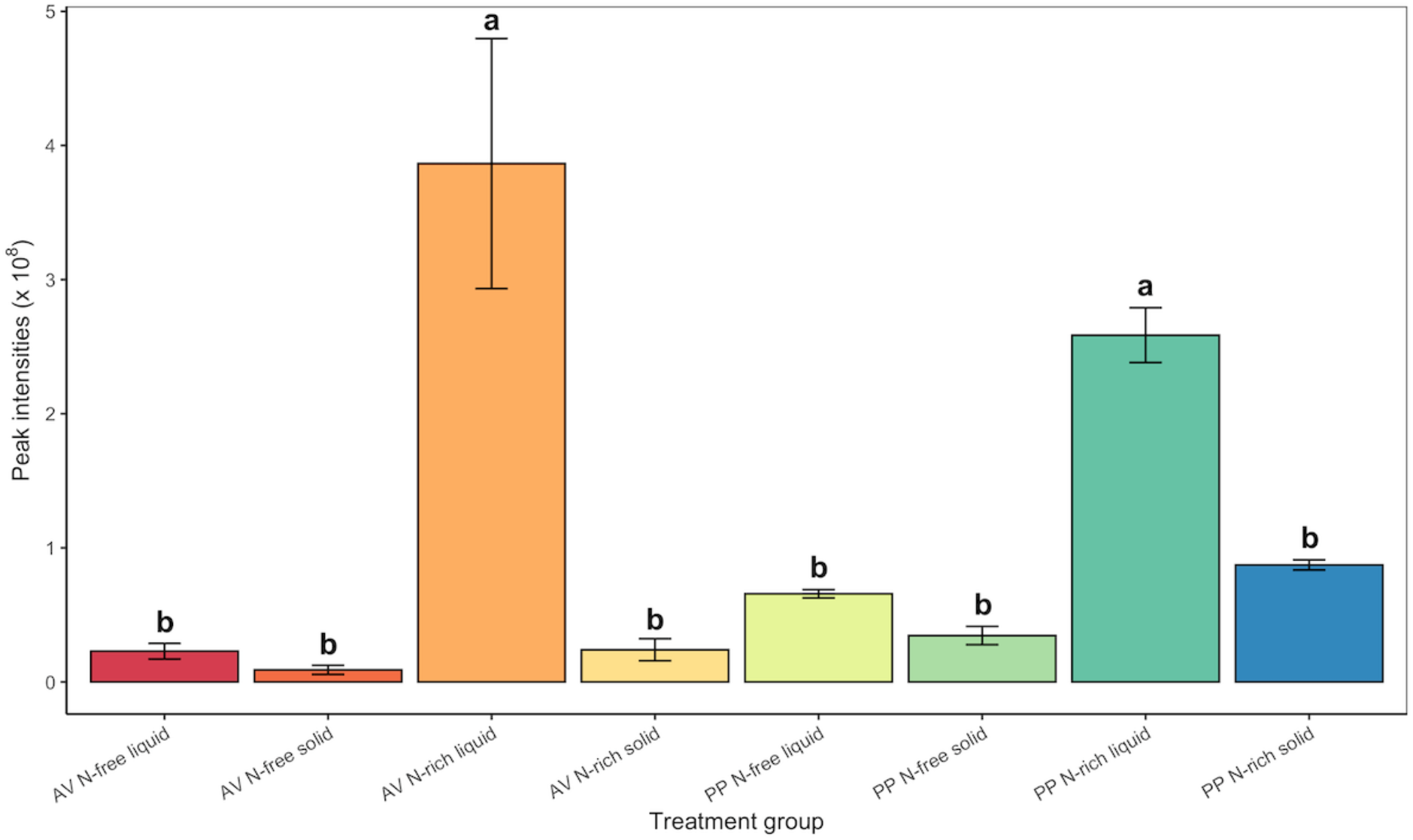
Peak intensities of N-containing metabolites detected across treatment groups. Bars represent average peak intensity ± standard error. Lowercase letters indicate significant difference at p < 0.05.

### Spatially resolved extracellular metabolites

Across all treatments, METASPACE analysis identified 69 metabolites in spatially resolved samples of which 41 were N-containing. However, only a few potential amino acids were detected at this resolved microbial scale including L-leucine and L-valine. These were only at detectable concentrations within the N-rich treatment (Fig. 7) unlike the diversity of amino acids detected in bulk samples predominately in association with N-free treatments. Observationally, N-free treatments seemed to be characterized by unique presence of organic acids rather than N-containing compounds. However, we did identify a few N-containing compounds unique to N-free treatments at the microbial scale including inosine and 4-pydroxic acid (Fig. 7). Inosine was detected in N-free treatments of both AV and PP and was not at detectable levels in N-rich treatments. Also, much like bulk sampling scale detection, 4-pyridoxic acid was exclusively detected in AV N-free treatment samples.

**Fig. 7:**
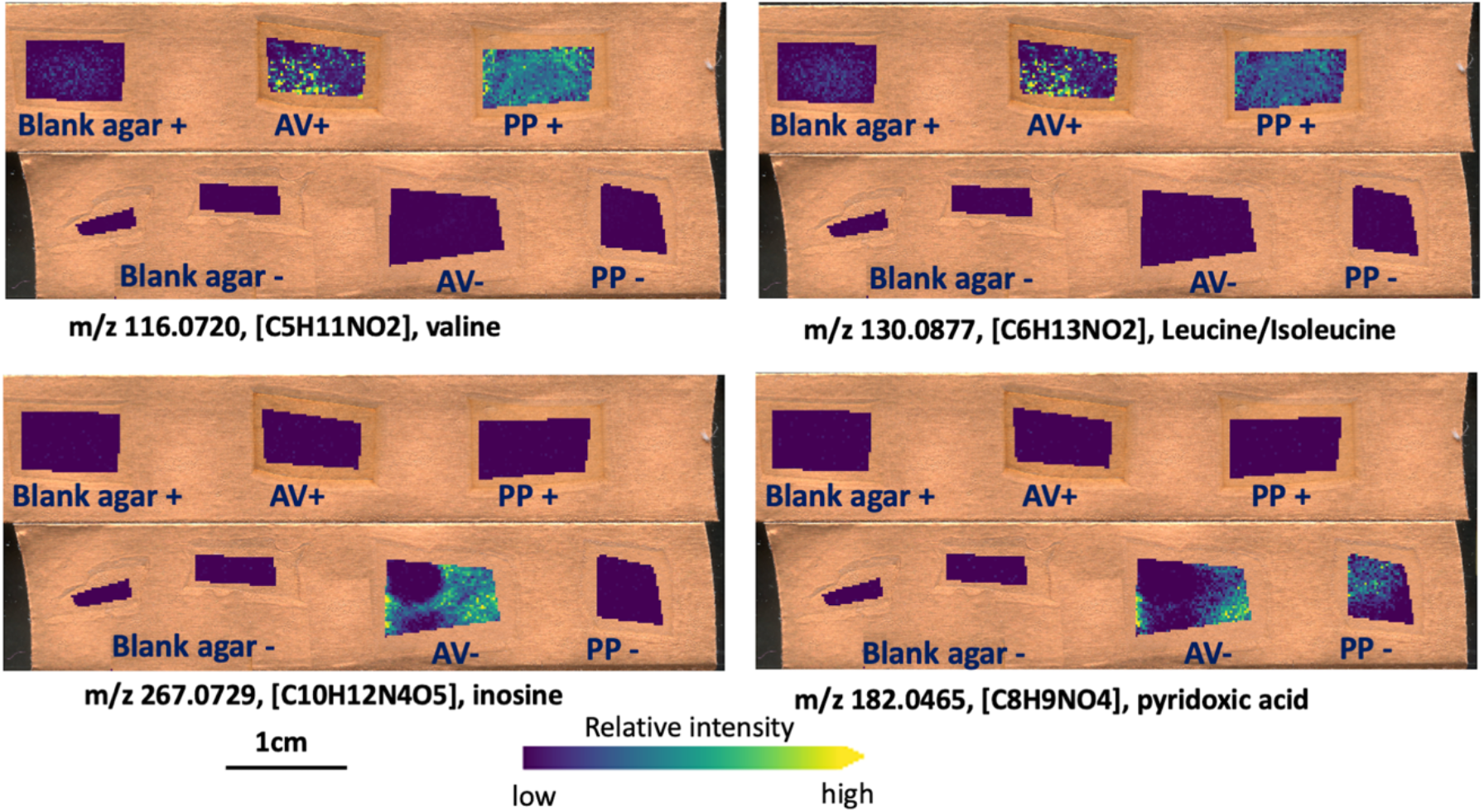
Examples of the N-containing metabolites detected at the microscale using MALDI MSI. All ions are annotated as [M–H]-adducts. Ion images of individual m/z values were generated on the same color bar scale for visual comparison in terms of relative ion abundance.

## Discussion

We explored the impact of N availability and physical structure on the extracellular metabolomics of diazotrophic bacteria across sampling scales from the spatially resolved to bulk. We find evidence of extracellular organic and inorganic N contributions from FLNF underscoring a key mechanism of terrestrial N contributions from FLNF. In general, we find physical structure and microbial function (e.g. FLNF) alter extracellular metabolite profiles and influence the detection of metabolites at bulk and spatially resolved scales.

### Nitrogen contributions from FLNF

Products of BNF by symbiotic diazotrophs are well-studied and typically observed as ammonia and ammonium with contested evidence for production of amino acids (18–21). This knowledge of symbiotic BNF is thought to translate directly to FLNF leading to the assumption that free-living diazotrophs also excrete ammonia/ammonium into the surrounding environment during BNF. However, ammonia produced during FLNF is rapidly assimilated through conversion to glutamine or glutamate via the glutamine synthetase (GS) and glutamate synthetase (GOGAT) pathways (34). Thus, excreted ammonium would necessarily be in excess of these assimilation pathways (34). Ammonium excretion has been observed in wild-type *Azotobacter vinelandii* DJ, at concentrations between ~2 and ~25 μM (35, 36), values within range of those measured in this study (Supp. Fig. 4). However, in many cases measurable ammonium excretion was only observed from *Azotobacter vinelandii* cultures genetically altered to disrupt the GS-GOGAT pathways or facilitate constitutive nitrogenase synthesis (37–40).

An alternative hypothesis to ammonium excretion is that N contributions occur as organic N, either through direct release of N-rich compounds like amino acids (18, 21) or through turnover of dead biomass (41). Our bulk metabolomics data support this hypothesis with many N-containing organic compounds, including amino acids detected in N-free treatments. In fact, N-free treatments were richer in N-containing compounds than N-rich conditions, particularly when comparing against N-rich solid media which had few N-containing metabolites (Supp. Fig. 3). The structure of our study did not allow us to determine whether these organic molecules were directly excreted by active, N-fixing cells or released during cell turnover. However, other metabolites detected in the system suggest cell turnover contributed at least partially to this N release. For example, we detected inosine in both bulk and spatially resolved analysis, and it was unique to N-free treatments in spatially resolved samples. Inosine, a metabolic product of adenine degradation likely indicates salvage activities by the bacterial populations (42, 43) and could indicate freely available nucleotides from cell lysis and turnover. FLNF may therefore contribute available N through increasing microbial biomass and turnover, but this needs to be verified in future studies. Regardless of whether these N-containing compounds are actively excreted or released after cell death, this metabolic exchange with the surrounding environment highlights a key mechanisms of terrestrial N contributions from FLNF.

### In vitro vs in situ analysis – identification of biosignatures

Through bulk and spatially resolved analysis, we found few N-containing metabolites exclusive to N-free treatments. At bulk scale, these include pantothenic acid, L-pyroglutamic acid, L-glutamic acid, and 4-pyridoxic acid. We similarly find 4-pyridoxic acid at the spatially resolved microbial scale as well as nine other metabolites including inosine. 4-pyridoxic acid was unique to AV N-free treatments at the microbial scale. However, despite being uniquely associated with N-free treatments and therefore microbial populations actively fixing N, it may be difficult to assign these as a signature of FLNF function. Of these compounds, only L-glutamic acid has a direct association with the FLNF pathway. Other metabolites seem more indicative of microbial nutrient needs and function. For example, pantothenic acid, vitamin B5, is involved in the synthesis of coenzyme A and is a coenzyme for many reactions involved in protein and lipid metabolism (44–46). This is particularly important for the processing of organic acids like malate, the main C source provided in this study. Thus, the detection of vitamin B5 is likely indicative of malate metabolism via the TCA cycle and its unique detection in the N-free treatment suggests a higher respiration rate in these N-fixing populations than in the N-rich populations. Increased respiration is a common response among diazotrophs in oxygenated environments as a protection mechanism to prevent or reduce denaturation of nitrogenase via oxygen (17, 47, 48). We also identified 4-pyridoxic acid, a derivative of pyridoxine (vitamin B6). Pyridoxine is a key cofactor in amino acid, fatty acid, and carbohydrate metabolisms, but can also act an oxygen protectant (46). During this redox reaction, pyridoxine degrades and can result in 4-pyridoxic acid. *A. vinelandii* has been observed to produce B vitamins while under diazotrophic conditions and this seems to be a hallmark of FLNF for this organism (46, 49, 50). Though not directly associated with the N-fixation pathway, these vitamins may tangentially indicate bacteria functions surrounding FLNF such as oxygen regulation and highlight the need to analyze bacterial function holistically rather than focusing on single reactions or pathways.

Additionally, the limited number of unique extracellular metabolites detected in N-free treatments suggests some microbial functions may not have detectable or unique biosignatures, in the form of extracellular metabolites. This is an important consideration when applying metabolomics to the study of complex soil systems. Soil metabolomics are increasingly being used to study soil microbial ecology and biogeochemical function and have been successfully applied to soil carbon cycling (51–54). However, metabolites are by definition the by-products of and substrates for metabolic function, and turnover rapidly in soils (55, 56). Therefore, typical soil extractions to collect extracellular components (e.g. K2SO4 extracts, leachate; (57, 58) only capture what is not consumed by the microbial community. This includes metabolites available in dissolved organic matter pools at the time of sampling and metabolites readily exchangeable from mineral surfaces (59). In both cases, metabolites could be temporally separated from their originating processes making it difficult to trace back the associated metabolic pathway. It could be even more challenging to capture metabolic biosignatures from nutrient-limited communities, such as those in bulk soil. Under nutrient-limited conditions, resulting metabolic products are likely to be rapidly assimilated or, in the case of processes like FLNF, not released to the surrounding environment. The potential signature compounds of FLNF found here (e.g. amino acids and B-vitamins) are also not uniquely produced by FLNF processes and would be difficult to directly link to FLNF *in situ*. We acknowledge our study system may provide a biased view on this issue, being a closed incubation system unlike soils where metabolites may diffuse away from microbes and persist in the environment. However, these findings highlight a key need to understand the soil microhabitat (11).

Lastly, our work highlights the importance of considering culture conditions and their association to *in situ* conditions as we observed clear differences in metabolite profiles between liquid and solid culture. Interestingly, though culture type strongly influenced metabolite profiles, it played a secondary role to N treatment in influencing the number of detected metabolites. This was particularly notable when N was readily available, where presence-absence based profiles were distinct between liquid and solid culture, but only under N-rich conditions. These differences in metabolite profiles were likely not driven by differences in biomass production as culture type had small and non-significant impacts on microbial metrics, like total biomass, and biomass C and N content. Thus, these responses seem specifically associated with the presence or absence of physical structure in the environment. Additionally, these findings suggest nutrient limitation, as experienced in the N-free treatments, may be a stronger driver of microbial activity than physical structure and simplified liquid culture may be somewhat informative to nutrient-limited *in situ* conditions.

### Implications scaling from microbial scale to bulk sampling

The combination of techniques in this study allowed us to explore detection of metabolites across scales from spatially resolved, relevant to microorganisms, to bulk, relevant for soil microbial ecology analysis. MALDI MSI allowed us to resolve abundances of extracellular metabolites on solid media at a microbial scale. Using GC-MS, we were able to evaluate detection of extracellular metabolites at the bulk scale. While the detection ranges of these two techniques do not fully overlap (50 – 500 m/z for GC-MS and 92 – 700 m/z for MALDI), many metabolites of interest to this study are measurable with both techniques providing valuable information about metabolite detection and sampling scale. It is also important to note that a lack of detection is not equivalent to metabolite absence but only indicates metabolite concentrations were below detection.

Through bulk sampling, we found a wide variety of N-containing compounds in N-free samples, but generally lower abundances of N-containing compounds than in the N-rich treatment. While N-containing compounds are characteristic of N-free samples at a bulk scale, these treatments had fewer N-containing metabolites when spatially resolved at the microbial scale. Interestingly, there was a shift in amino acid detection between spatially resolved and bulk scales where amino acids were commonly detected in N-free samples at bulk scale, but in N-rich samples when spatially resolved. This somewhat counterintuitive result highlights differences in N competition at the microbial scale and its influence on bulk measurements.

First, detection of a diverse array of amino acids in the N-free treatment in bulk sample, but not in spatially resolved samples suggests N competition at the microbial scale resulted in rapid uptake of amino acids, while extraction of the bulk metabolite pool likely captured the cumulative low abundance signal of the entire system. Amino acids are shown to have short residence times in soils and experience rapid uptake and turnover (60, 61). In the case of microbial vs. bulk scale, it is likely the spatially resolved pool of extracellular amino acids collected from microbial colonies (~200 μm spatial resolution) was small and often below detection. However, in bulk sampling of millions of cells, a larger pool of amino acids coupled to our sampling method could have allowed amino acids to diffuse away and accumulate to detectable levels.

Second, biofilm formation is likely to influence diffusion of metabolites into the surrounding environment (62, 63). Bacteria tend to live in biofilms in their natural environments rather than as individually dispersed cells (64). However, the impact of surrounding environmental conditions, including nutrient availability, on biofilm production is unclear. For example, some studies suggest nutrient limiting conditions may promote greater biofilm formation (65), while others suggest biofilm formation is greater under more favorable growth conditions (66, 67). This is particularly notable for diazotrophs as biofilms can play a role in oxygen protection (47), thus investment in biofilm could be beneficial to FLNF activity. Yet, under severe N limitation imposed by an N-free environment the high energy demands of FLNF may limit investment in biofilm. While not directly measured in this study, we noted solid agar plates of *Azotobacter vinelandii* and *Paenibacillus polymyxa* cultures had visually greater biofilm production under N-rich than N-free conditions. Thus, diffusion of amino acids away from populations would have been more easily achieved in the N-free treatment. This is evidenced by the similarity between metabolite chemistry in liquid and solid culture under N-free treatments (Fig. 5B). Similarly, a small number of amino acids were detected in spatially resolved samples from N-rich treatments, but not in bulk samples for the similar N-rich solid media treatment. This could have resulted from greater biofilm formation under N-rich conditions and limited diffusion of small molecules away from cell populations.

The detection of small molecules across scales has important implications for the influence of soil microbial communities on their surrounding environment. In general, our results indicate microbial scale processes drive bulk metabolite availability. The N-rich treatment in this experiment is an optimal environment and most representative of carbon and nutrient rich soil environments like the rhizosphere or detritusphere. Our findings suggest these conditions would result in production of valuable small molecules, like N-rich amino acids, potentially exchangeable with the immediate environment, but biofilm formation may limit diffusion far into the soil environment. Under limiting conditions of the N-free treatment, similar those of bulk soil, microbial activity produces valuable metabolites, like amino acids, but competition between microbes reduces the exchange of these molecules. Understanding how these differences in microbial scale conditions influence microbial activity and detectability of function is crucial to accurately linking microbe and ecosystem.

## Conclusions

We demonstrate extracellular production of inorganic and organic N during FLNF and reveal the importance of habitat conditions and sampling scale when quantifying microbial activity. Across bulk and spatially resolved sampling scales, we identified metabolites uniquely associated with FLNF activity including several B-vitamins, which may play roles in mitigating oxygen damage to nitrogenase. Despite finding unique metabolites and potential biosignatures, many detected metabolites are not exclusively produced through FLNF related pathways, thus would be difficult to assign to FLNF for *in situ* soil samples. This would likely hold true for other processes under nutrient limited conditions where metabolic products are rapidly assimilated and not captured during sampling. Our findings highlight the need to carefully consider both structural complexity and sampling scale when quantifying microbial function. We found culture conditions to be a key driver of metabolite chemistry under N-rich and N-free conditions, indicating physical structure influences microbial processes. Across scales, our results indicate high N competition at the microbial scale under N-free conditions, while at the bulk scale N appeared readily available within the microbial environment. These differences in environmental conditions across scales could lead to incorrect interpretations of microbial function as immediate conditions surrounding microorganisms will drive their activity and may not necessarily match what is measured through bulk or composite sampling.

## Materials and Methods

### Culture conditions

Two diazotrophic bacteria with distinct growth strategies (e.g. gram-negative vs gram-positive) and fully sequenced genomes (22, 23)were chosen for this study, *Azotobacter vinelandii* DJ (ATCC BAA 1303; hereafter AV) and *Paenibacillus polymyxa* (ATCC 842; hereafter PP). Bacteria were cultured under N-free and N-rich conditions, respectively promoting or inhibiting FLNF. Nfb media, commonly used to isolate diazotrophs (24), was used for N-free treatments, and was supplemented with tryptone for N-rich treatments. Both treatments contained 1.79 g C L^−1^ as malic acid and N-rich media contained ~1.33 g N L^−1^ as tryptone. Cultures were grown in liquid or solid agar media and all media was autoclave sterilized prior to inoculation.

Thirty samples were cultured (2 organisms × 2 N treatments × 2 media types × 3 replicates, plus controls) for bulk analysis with an additional set of 14 samples (2 organisms × 2 N treatments × 3 replicates, plus controls) on solid media for spatially resolved analysis. Cultures were grown in a temperature-controlled incubator at 25°C to 10^7^ CFU mL^−1^, based on liquid cultures, and then harvested for analysis of extracellular metabolites at two scales – bulk sampling via MPLEx extraction and GC-MS (25) and spatially-resolved sampling via colony analysis with MALDI MSI. Extracellular ammonium availability and microbial biomass, including total biomass, biomass carbon (C) and biomass N, were also measured. Because FLNF activity is necessary for microbial growth under N-free conditions, measures of total biomass and biomass N are used as estimates of FLNF (24).

### Sample collection

Extracellular metabolites were collected from liquid culture by centrifuging culture tubes to pellet cells and collecting the resulting supernatant for bulk analysis as described below. Cell pellets were resuspended in autoclave sterilized nanopure water, immediately flash frozen on liquid nitrogen, and stored at −80°C until further analysis. Extracellular metabolites were collected from solid media for bulk and spatially resolved analysis. Bulk samples were collected by washing culture plate surfaces with autoclave sterilized nanopure water and collecting the resulting rinsate. Samples were collected for spatially resolved analysis as described below. Lastly, microbial colonies from rinsate plates were collected from the surface by gentle scraping, transferred to autoclave sterilized nanopure water, flash frozen on liquid nitrogen, and stored at −80°C until further processing.

### Microbial biomass – total biomass, biomass C, and biomass N

Frozen cell pellets and colonies were lyophilized until completely dry and weighed to obtain total biomass, including cells and associated debris such as exopolysaccharides (EPS). Dried biomass was ground using sterile steel beads and then analyzed for C and N content on a VarioTOC Cube (Elementar, Langenselbold, Germany).

### Extracellular ammonium availability

We measured extracellular ammonium concentrations in supernatant and rinsate samples using a high-throughput colorimetric ammonium assay (26). Briefly, samples were pipetted in triplicate into clear 96-well plates and incubated with ammonium salicylate and ammonium cyanurate reagents to facilitate color change via the Berthelot reaction. Plates were read for absorbance at 610 nm on a Synergy H1 plate reader (BioTek Instruments, Inc., Winooski, VT, USA).

### Bulk metabolomics – GC-MS

Bulk metabolomics analysis was conducted on 1 ml subsamples of undiluted, supernatant and rinsate samples. Supernatant and rinsate samples were prepared for metabolite analysis via GC-MS following the MPLEx protocol for simultaneous metabolite, protein, and lipid extraction (25). Additionally, 1 ml of supernatant and rinsate from sterile liquid culture and solid culture plates were also extracted via MPLEx as background controls. This extraction method allows simultaneously collection of metabolites, lipids, and proteins; however, lipid fractions were not analyzed in this study. Additionally, protein yields were too low for downstream analysis. Metabolite samples were completely dried under speed-vacuum concentrator and chemically derivatized prior to analysis by GC-MS as reported previously (27). The m/z range of derivatized metabolites scanned was 50 - 550 m/z which can detect organic acids, amino acids, and mono to tri-saccharides. Raw GC-MS data were processed using the PNNL in-house metabolomics database, which can identify metabolites using two dimensional matching factors (fragmented spectrum + retention index (28), and with cross-checking against commercially available NIST 20/Wiley 11^th^ GC-MS spectral databases (25, 29).

### Spatially resolved metabolomics – MALDI MSI

Samples were prepared for spatially resolved analysis via MALDI-MSI using a previously described workflow (30). Briefly, areas of agar were excised from Petri dishes and placed onto double-sided adhesive copper tape adhered to indium tin oxide (ITO)-coated glass slides (Bruker Daltonics; Supp. Fig 1.). This approach enhanced our sensitivity for analysis in negative ionization mode and improved adhesion of agar onto the MALDI target. Samples were dried at room temperature overnight, then treated with MALDI matrix using a HTX TM-Sprayer (HTX Technologies). For analysis in negative-ion mode, 7 mg mL^−1^ of N-(1-naphthyl) ethylenediamine dihydrochloride (NEDC) in 70% MeOH was sprayed with eight passes at 1,200 μL min^−1^, 75°C, a spray spacing of 3 mm, and a spray velocity of 1,200 mm min^−1^. MALDI-MSI was performed on a 15-Tesla Fourier transform ion cyclotron resonance (FTICR)-MS (Bruker Daltonics, Billerica, MA, USA) equipped with SmartBeam II laser source (355 nm) using 200 shots pixel^−1^ with a frequency of 2 kHz and a step size of 200 μm. FTICR-MS was operated to collect m/z 92–700, using a 209-ms transient, which translated to a mass resolution of R ~ 70,000 at 400 m/z. Metabolites in this range can typically be detected to fmol concentrations.

### Data Analysis

A factorial ANOVA with N treatment, culture type, organism and their interactions as main effects followed by a Tukey’s post hoc test was used to determine treatment differences for measured variables. Prior to statistical analysis, bulk metabolite values were blank corrected by subtracting peak intensities identified in background controls of the associated treatment. Differences in bulk metabolite chemistry were evaluated using distance matrices based on peak intensities (Bray-Curtis) presence-absence (Jaccard) generated from all detected metabolites using R *vegan* (31). Differences between culture type, N treatment, and organism were determined via PERMANOVA using *adonis* in R *vegan*. Spatially resolved metabolite data was acquired using FlexImaging (v 4.1, Bruker Daltonics), and image processing, segmentation, co-localization analysis and visualization were performed using SCiLS (Bruker Daltonics). The list of m/z values that colocalized with the colonies were uploaded to the METLIN (https://metlin.scripps.edu) for putative molecular annotations based only on accurate m/z, secured by using a 3-ppm window during the search. imzML files (created by SCiLS) of our analyses were also uploaded to METASPACE (32) for metabolite annotation based on both accurate m/z and a comprehensive bioinformatics framework that considers the relative intensities and spatial colocalization of isotopic peaks as well as quantifies spatial information with a measure of spatial chaos followed by the estimation of the False Discovery Rate. For this purpose, we used KEGG-v1 and NPA-2019-08 (Natural Product Atlas) databases that are available in METASPACE. METASPACE uses by default 3 ppm window in its annotation engine.

## Acknowledgements

We would like to thank Heather Olson, Carrie Nicora, and Jesse Trejo for sampling processing work and for taking time to carefully train D.N.S. on the methods. D.N.S. is also grateful for the support of the Linus Pauling Distinguished Postdoctoral Fellowship program through Pacific Northwest National Laboratory. Pacific Northwest National Laboratory is a multiprogram national laboratory operated by Battelle for the U.S. Department of Energy under contract DE-AC05-76RL01830. This research was performed using resources at the Environmental Molecular Sciences Laboratory (EMSL; grid.436923.9), a DOE Office of Science User Facility sponsored by the Biological and Environmental Research program.

