## Supplemental material for "Bulk and spatially resolved extracellular metabolomics of free-living nitrogen fixation"

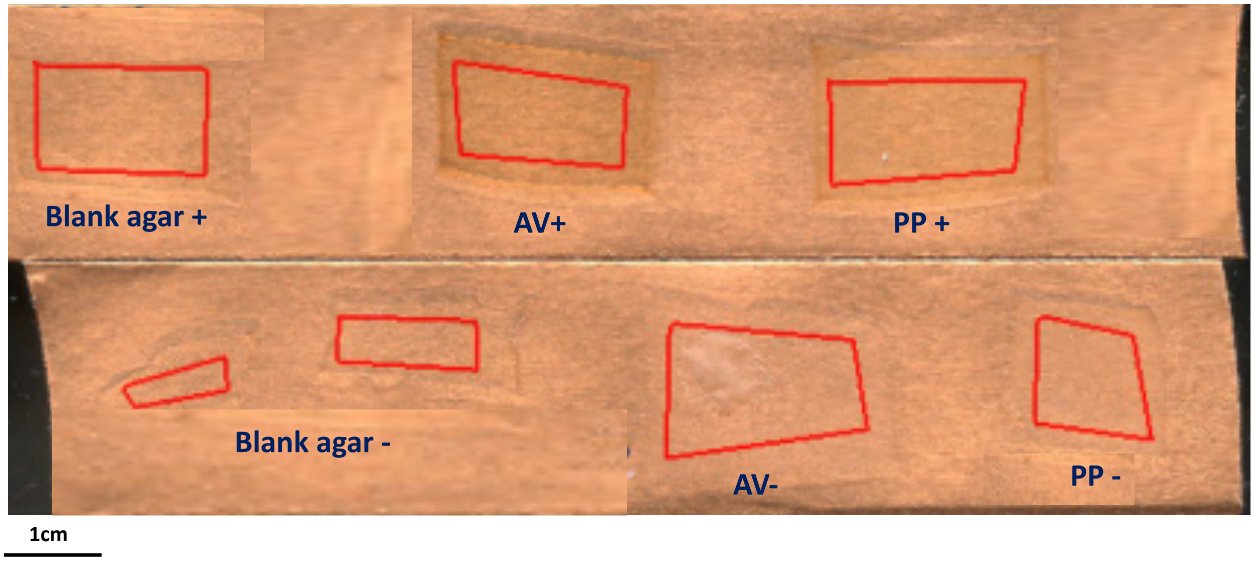


**Supp. Fig. 1:** Optical image of agar areas collected for MALDI MSI. Areas are placed on double-sided adhesive copper tape and adhered to indium tin oxide coated glass slides. Images were taken after regions were coated with MALDI matrix application. Red regions denote MALDI target areas within each sample.

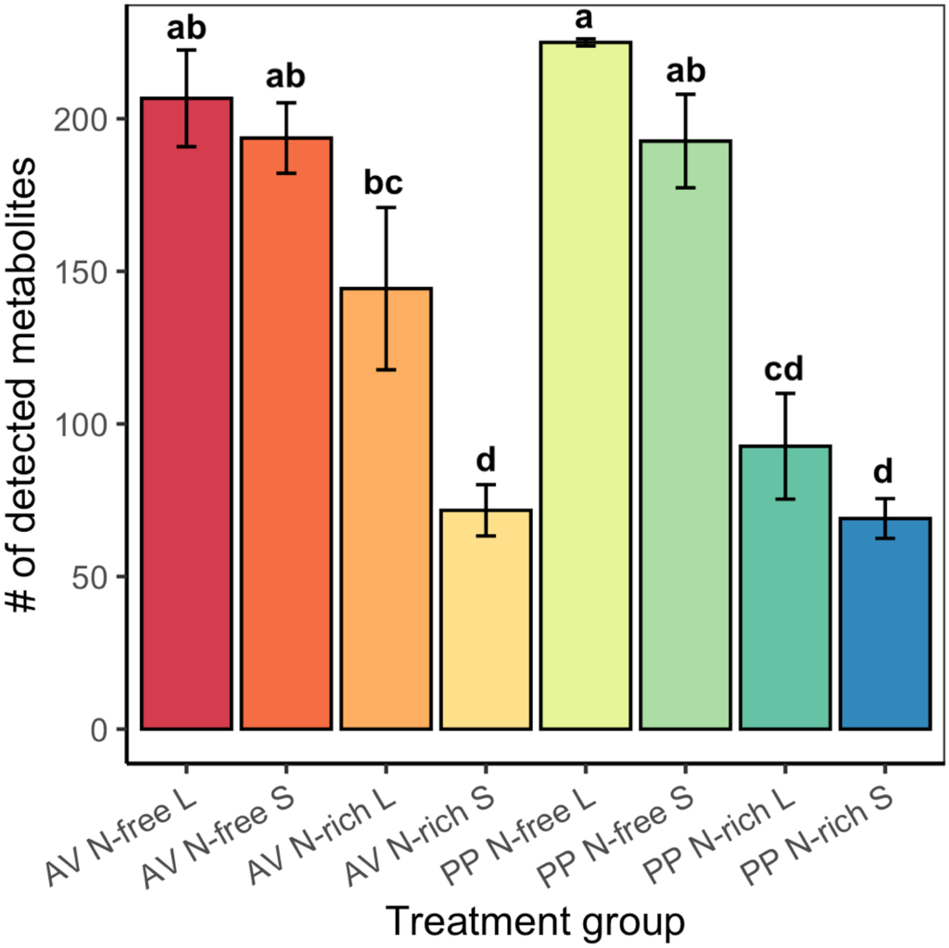


**Supp. Fig. 2:** Number of detected metabolites at the macroscale across treatments. Bars represent average ± standard error. Lowercase letters indicate significant difference at $p < 0.05$.

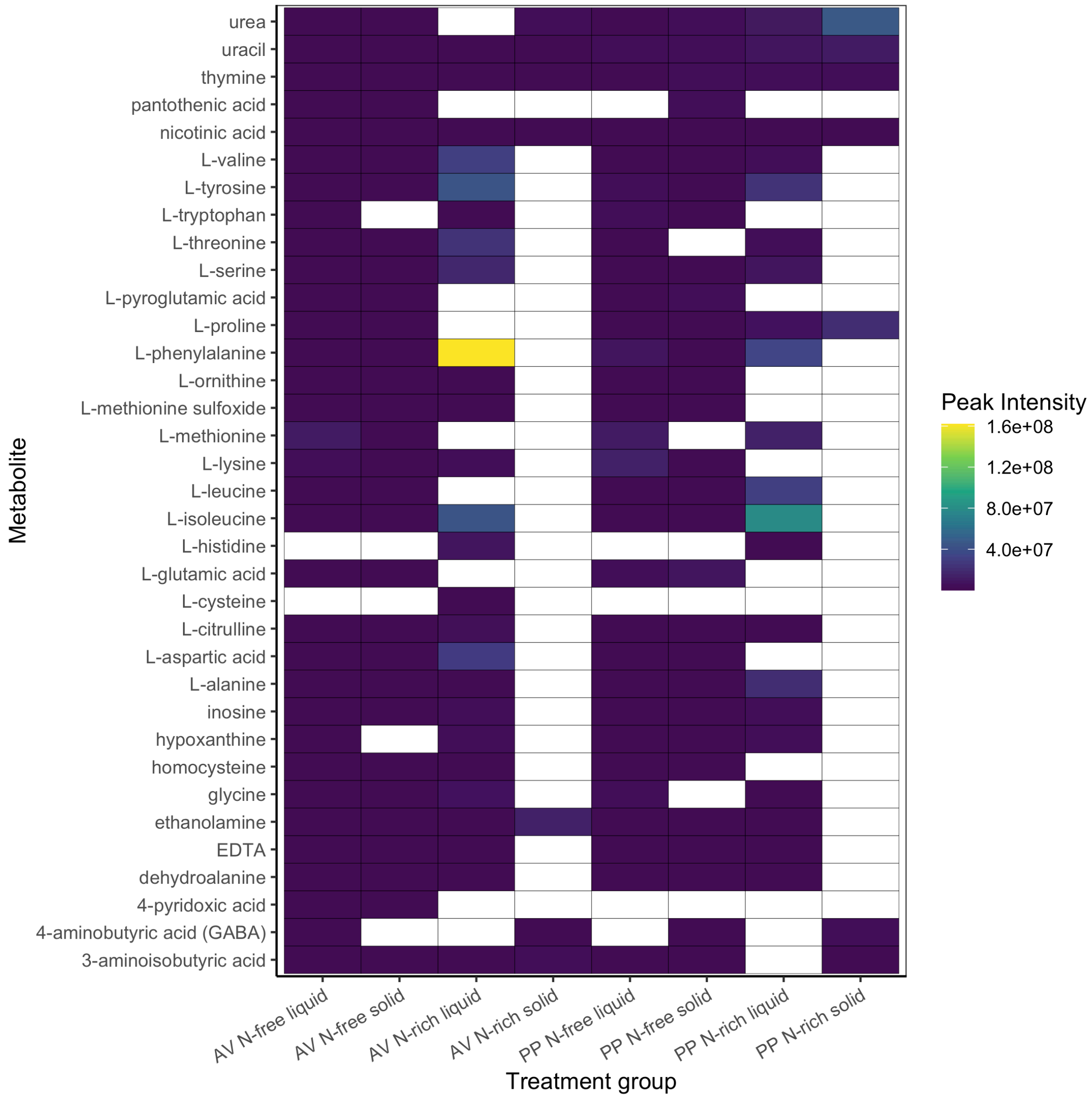


**Supp. Fig 3**: Heatmap of N-containing metabolites and their peak intensities across treatments. White cells indicate metabolite abundance was below detection.

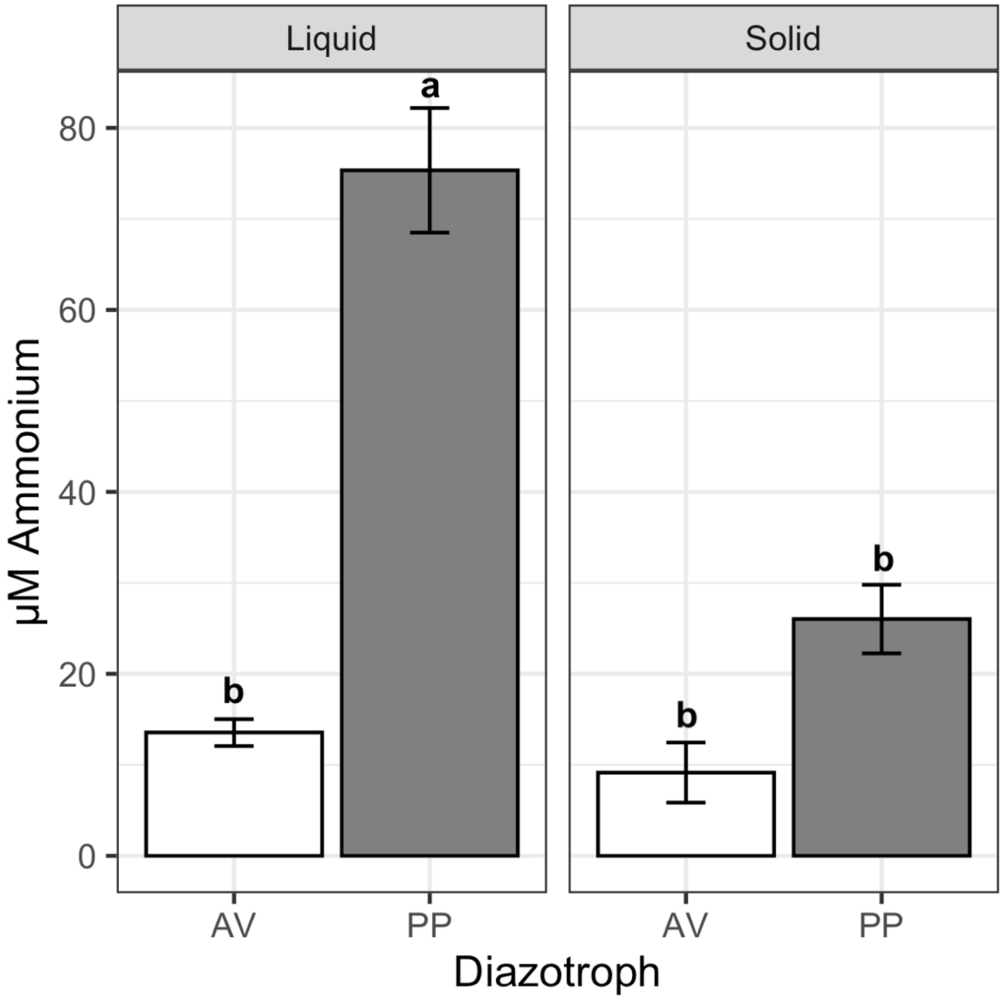


**Supp. Fig 4**: Extracellular ammonium availability in µM for N-free treatments. Bars represent average values ± standard error. Lowercase letters indicate significant difference within culture type (liquid or solid) at $p < 0.05$.
